# An aminoacylation independent activity of PheRS/FARS promotes growth and proliferation

**DOI:** 10.1101/2020.10.29.360420

**Authors:** Manh Tin Ho, Jiongming Lu, Beat Suter

## Abstract

Aminoacyl-tRNA synthetases (aaRSs) not only load the appropriate amino acid onto their cognate tRNA, but many of them perform additional functions that are not necessarily related to their canonical activities. Phenylalanyl-tRNA synthetase (PheRS/FARS) levels are elevated in various cancer cells compared to their normal cell counterparts. However, whether and how these levels might contribute to tumor formation was not clear. Here, we show that PheRS is required for cell growth and proliferation. Interestingly, elevated expression of the α-PheRS subunit alone stimulates cell growth and proliferation. In the wing discs system, this leads to a strong increase of mitotic cells. Clonal analysis of twin spots in dividing follicle cells revealed that elevated expression of the *α-PheRS* subunit causes cells to grow and proliferate about 25% faster than their normal twin cells. Importantly, this stimulation of growth and proliferation neither required the β-PheRS subunit nor the aminoacylation activity, and it did not visibly stimulate translation. These results, therefore, revealed a non-canonical function of an ancient housekeeping enzyme, providing novel insight into its roles in health and diseases.

## Introduction

Many cancer tissues display higher levels of Phenylalanyl-tRNA synthetase (PheRS, FARS, or FRS) than their healthy counterparts according to the database “Gene Expression across Normal and Tumor tissue” (GENT2) (Park et al., 2019). Interestingly, a correlation between tumorigenic events and PheRS expression levels had been noted already much earlier for the development of myeloid leukemia (Sen et al., 1997) and this was further supported by additional results from the Safro group (Rodova et al., 1999). Despite this, a possible causative connection between elevated PheRS levels and tumor formation had so far not been reported and, to our knowledge, also not been studied. This might have been due to the assumption that higher PheRS levels could simply reflect the demand of tumorigenic cells for higher levels of translation, or it could have to do with the difficulty of studying the moonlighting function of a protein that is essential in every cell for basic cellular functions such as translation.

Aminoacyl-tRNA synthetases (aaRSs) are a group of enzymes that charge transfer RNAs (tRNAs) with their cognate amino acid, a process called aminoacylation which serves as a key step for protein translation. This activity makes them essential for the accurate translation of the genetic information into polypeptide chains (Schimmel and Soll, 1979). Besides their canonical role in translation, an increasing number of aaRSs have been shown to perform additional functions in the cytoplasm, the nucleus, or even outside of the cell (Casas-Tinto et al., 2015; Gomard-Mennesson et al., 2007; Greenberg et al., 2008; Guo and Schimmel, 2013; Lee et al., 2004; Nathanson and Deutscher, 2000; Otani et al., 2002; Smirnova et al., 2012; Zhou et al., 2014). For example, the amino-acid binding site of LysRS has an immune response activity; or TrpRS inhibits the vascular endothelial (VE)-cadherin and through this elicits an anti-angiogenesis activity (Tzima et al., 2005; Yannay-Cohen et al., 2009).

Cytoplasmic PheRS, a heterotetrameric protein consisting of 2 alpha- (α) and 2 beta-(β) subunits responsible for charging tRNA^Phe^ with phenylalanine is one of the most complex members of the aaRSs family (Roy and Ibba, 2006). The α subunit includes the catalytic core of the tRNA synthetase and the β subunit contains structural modules with a wide range of functions, including tRNA anticodon binding, hydrolyzing mis-activated amino acids, and editing wrongfully aminoacylated tRNA^Phe^ species (Ling et al., 2007; Lu et al., 2014; Roy and Ibba, 2006). Importantly, both subunits are required for the aminoacylation of tRNA^Phe^.

We set out to address the question of whether and how elevated levels of PheRS can contribute to over-proliferation. To test for this activity, we studied the role of PheRS levels in the Drosophila model system with the goal of finding out whether elevated levels of PheRS allow higher translation activity or whether a moonlighting role of *PheRS* might provide an activity that might contribute to elevated growth and proliferation. We found that α-PheRS levels regulate cell growth and proliferation in different tissues and cell types. Interestingly, elevated levels of α-PheRS do not simply allow higher levels of translation. Instead, α-PheRS performs a moonlighting function by promoting growth and proliferation independent of the β-PheRS subunit and even if it lacks the aminoacylation activity.

## Results

### PheRS is required for growth and proliferation

The *Drosophila* FARSA and FARSB homologs *α-PheRS* and *β-PheRS* are essential genes (Lu et al., 2014). To find out whether cellular levels of PheRS might correlate with and possibly contribute to growth, we tested whether reduced levels in specific tissues affect the growth of the organ and animal. For this, we used RNAi to reduce their activity in two specific tissues, the eye, an organ that is not essential for viability, and the fat body (Fig 1A,B). Indeed, knocking down either of the two subunits in the developing eye dramatically reduced the size of the adult eye (Fig 1A). Similarly, reducing *α-PheRS* or *β-PheRS* expression levels in the larval fat body also caused a growth reduction. However, presumably because of its role in systemic growth (Texada et al., 2020), the fat body knockdown of *PheRS* reduced the size of the entire pupae (Fig 1B).

**Fig 1:**
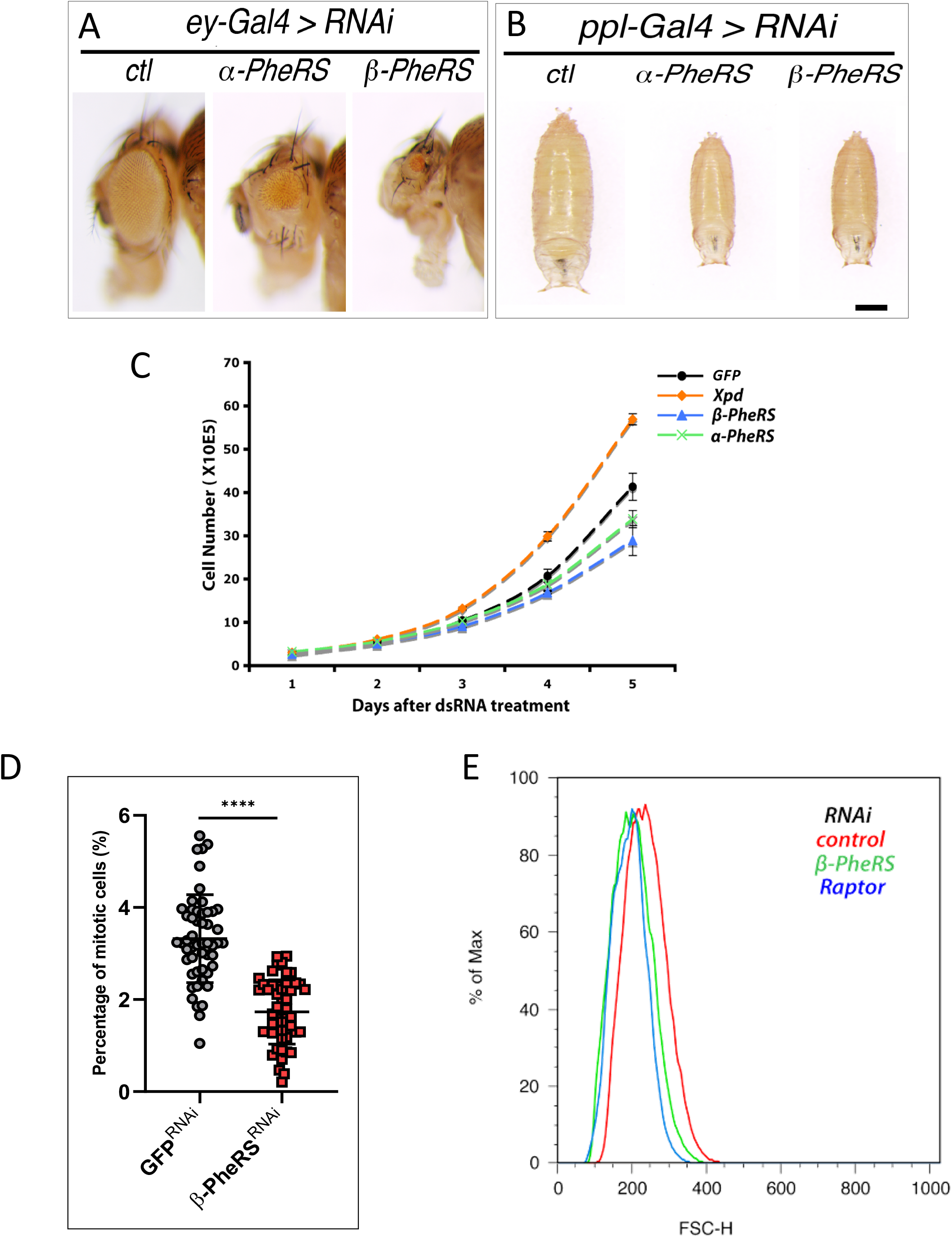
*PheRS* knockdown reduces growth. (A) *PheRS* knockdown in eye discs results in smaller eyes. *eyeless-Gal4* was used to drive the knockdown of the α and β-subunits (α-sub and β-sub). Control (ctl) is the driver alone. RNAi lines were obtained from VDRC. Scale bar represents 250 μm. (B) *PheRS* knockdown in the larval fat body results in smaller pupae. *ppl-Gal4* was used to drive the knockdown. The same RNAi lines were used as in (A). Scale bar represents 500 μm. (C) *PheRS* knockdown in Kc cells downregulates cell proliferation. Knockdown was performed by adding dsRNA to the cell medium. Cell numbers were determined during the days following the dsRNA treatment. The result represents two independent experiments. The negative control was GFP RNAi while the positive control was *xpd* RNAi. (D) *β-PheRS* knockdown in Kc cells reduces the frequency of mitotic cells. The mitotic index was determined based on the frequency of the phospho-Histone H3 positive cells. Over 10,000 cells were counted for each treatment (P<0.001). (E) *β-PheRS* knockdown in Kc cells decreases cell size. The forward scatter (FSC) was used to determine cell size. *Raptor* RNAi is a positive control, and *β-PheRS* knockdown showed similar cell size.

To further analyze the changes at the cellular level, the effect of knocking down *α-PheRS* and *β-PheRS* in *Drosophila* Kc cells was first examined at the level of cell proliferation (Fig 1C). The knockdowns were carried out by adding dsRNA into the medium, and the cell numbers were recorded during the following days. Compared to the controls, cells treated with *β-PheRS* RNAi started to show lower cell numbers on day 3, and the cell count was around 75% of the control on day 5. In Kc cells, knocking down either subunit alone reduces levels of the α- and the β-PheRS subunit (Lu et al., 2014). It was therefore reassuring that *α-PheRS* knockdown showed similar results. The lower number was not due to cell death, as judged by trypan blue staining of the cells before counting. No obvious increase in dead cells was detected upon RNAi treatment. As a positive control, *xpd* RNAi was performed and showed the published increased in cell proliferation (Chen et al., 2003b). The fact that knockdown of *xpd* can speed up cell growth and proliferation not only indicates that the Kc cells were healthy, but also that the PheRS levels are not limiting for growth and proliferation, but can sustain even higher proliferative activity.

Determining the mitotic index upon PheRS knockdown revealed that the reduced PheRS levels caused a strong reduction of mitotic cells as indicated by the lower fraction of phospho-Histone 3 (PH3; mitotic marker) positive cells. The RNAi treatment reduced the mitotic index to 1.7% ± 0.16, which corresponded to half of the control (3.3% ± 0.16) (Fig. 1D). Similarly, cell size was also affected by *PheRS* knockdown (Fig 1E). The cell size showed a reduction that was similar to the one observed upon RNAi-mediated knockdown of *raptor*, a component of the TORC1 signaling that regulates cell growth (Kim et al., 2002). These experiments showed that *α-PheRS* and *β-PheRS* are required for normal growth and proliferation of cells, organs, and entire animals. The results might reflect the requirement for the enzymatic activity of PheRS in charging the tRNA^Phe^ with its cognate amino acid phenylalanine, or it might point to a novel, possibly moonlighting function of PheRS in stimulating growth and proliferation.

### PheRS lacks apparent amino acid sensor activity for TORC1

The TORC1 signaling pathway activates growth and proliferation of cells depending on the availability of amino acids, growth factors, and energy (Laplante and Sabatini, 2012; Wullschleger et al., 2006). In addition to its canonical function in charging tRNA^Leu^ with leucine, the LeuRS serves as the central amino acid sensor in this pathway (Bonfils et al., 2012b; Han et al., 2012b). We, therefore, tested whether PheRS might also be involved in nutrient sensing for TORC1 signaling in an analogous way. Amino acid deprivation causes downregulation of phosphorylation of dS6K in Kc cells, and subsequent stimulation with amino acids restores phospho-dS6K levels (Kim et al., 2008). The Rag complex was identified as a nutrient sensor in this pathway, and knockdown of *RagA* prevents the TORC1 complex from sensing the availability of amino acids (Kim et al., 2008; Sancak et al., 2008). We, therefore, used *RagA* as our control (Fig 2A,B). In contrast to *RagA*, knocking down *β-PheRS* (which also reduces α-PheRS levels) did not prevent amino acids sensing in this assay and phosphorylation of dS6K was still induced to a similar level as in the control when amino acids or phenylalanine were re-added after deprivation (Fig 2A,B). In this case, it did not matter whether we added back all amino acids or only L-Phe. Although we cannot rule out that the RNAi knockdown was insufficient to reveal an amino acid-sensing function for PheRS, the results seem to suggest that PheRS might not serve as an aa sensor upstream of the TORC1 complex.

**Fig 2:**
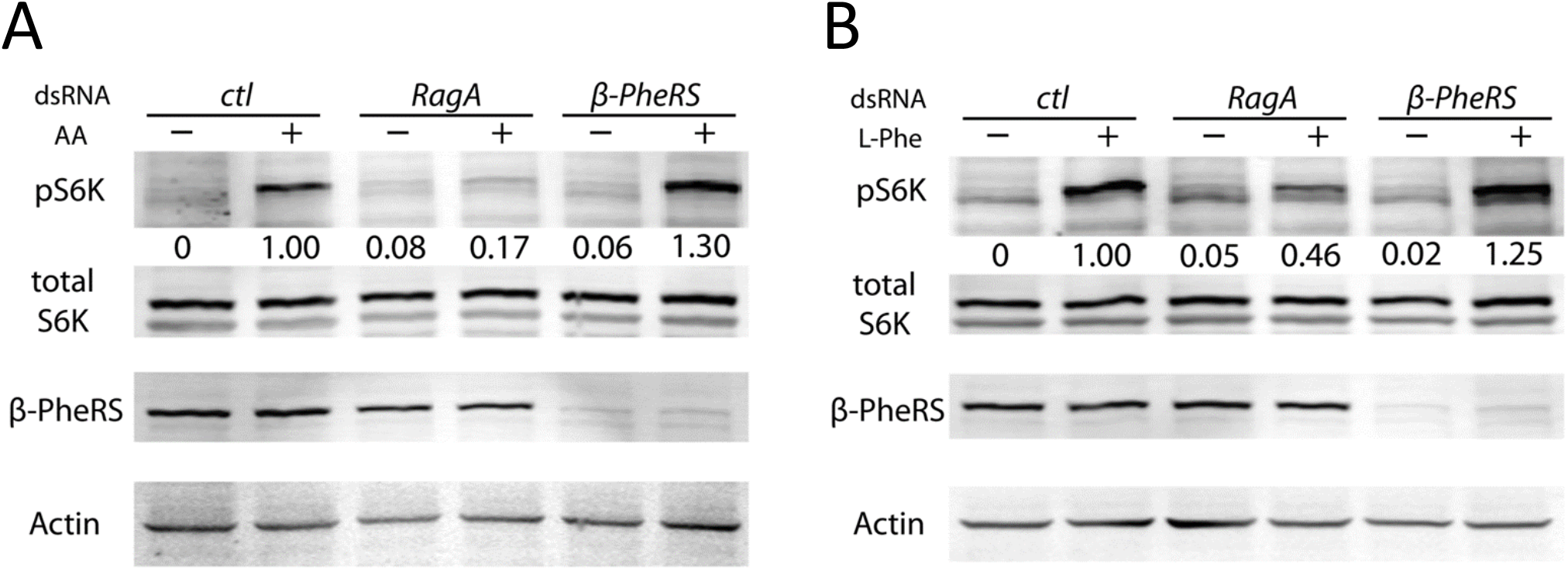
PheRS does not act as an amino acid sensor for the TORC1 complex. (A) *β-PheRS* knockdown cannot block the TORC1 complex from sensing the availability of amino acids. Phospho-S6K was used as a readout of TORC1 signaling. Starvation (-) was performed by depriving cells of amino acids for 30 mins, and stimulation (+) was performed by adding back amino acids for 30 mins after starvation. Control is a mock RNAi, *Rag A* RNAi is known to block the sensing of amino acids. Phospho-S6K levels were quantified relative to the Actin levels in the same extract. (B) *β-PheRS* knockdown did not block the TORC1 complex from sensing the availability of L-Phe. The same experiment as in (A) was performed but using L-Phe and L-Glu for stimulation. L-Glu was reported to be necessary for amino acid transport (Nicklin et al., 2009). Levels of phospho-S6K were quantified relative to the Actin levels in the same extract.

### α-PheRS activity is sufficient to induce additional M-phase cells

Circumstantial evidence suggests that elevated PheRS levels do not simply allow a higher translational activity to overcome a growth rate restriction imposed by hypothetically limiting levels of PheRS. PheRS is unlikely to be rate-limiting for cellular growth because animals with only one copy of *α-PheRS* (*α-PheRS*/-) or *β-PheRS* (*β-PheRS*/-) do not show a phenotype (Lu et al., 2014). Furthermore, Kc tissue culture cells can be stimulated to grow more rapidly without artificially over-expressing *α-PheRS* or *β-PheRS* (Fig 1C; (Chen et al., 2003a). To test whether elevated levels of PheRS can stimulate growth or proliferation, we expressed *α-PheRS*, *β-PheRS*, and both subunits together in the posterior compartment (P) of wing discs using a Gal4 driver under the control of the *engrailed*(*en*) promoter. This *en-Gal4* transcription factor drives the expression of the *α-PheRS* and *β-PheRS* open reading frames (ORFs) that were cloned behind a UAS promoter and integrated into a defined landing platform in the fly genome that also contains the endogenous *α-PheRS* and *β-PheRS* genes. In this assay, the anterior compartment expresses normal endogenous PheRS levels and serves as an internal control. The posterior compartment expresses the endogenous *α-PheRS* and *β-PheRS* genes and the two transgenes driven by *en-Gal4*. When α-PheRS and β-PheRS levels were raised in the posterior wing disc compartment together, the PH3 labeling revealed a 40% increase in mitotic cells in the posterior (P) compartment relative to the anterior (A) one of the same wing disc (Fig 3D). Surprisingly, the same result was obtained when only the levels of the α-PheRS subunit alone were raised (Fig 3A-D), but not when only the β-PheRS subunit levels were raised (Fig 3D). In addition, raising the α-PheRS subunit levels alone did not affect the β-PheRS subunit levels (Fig 3C-C”’). These results pointed to the possibility that this cell cycle effect might not only depend on the α-PheRS subunit alone but that it might also be a translation independent function of α-PheRS because aminoacylation requires both PheRS subunits to ligate Phe to the tRNA^Phe^ (Lu et al., 2014; Mosyak et al., 1995); Fig 4A). The fact that elevated α-PheRS levels alone are sufficient to raise the mitotic index is therefore another strong indication that α-PheRS possesses a proliferative activity that is unlikely to be mediated by increased tRNA^Phe^ aminoacylation.

**Figure 3:**
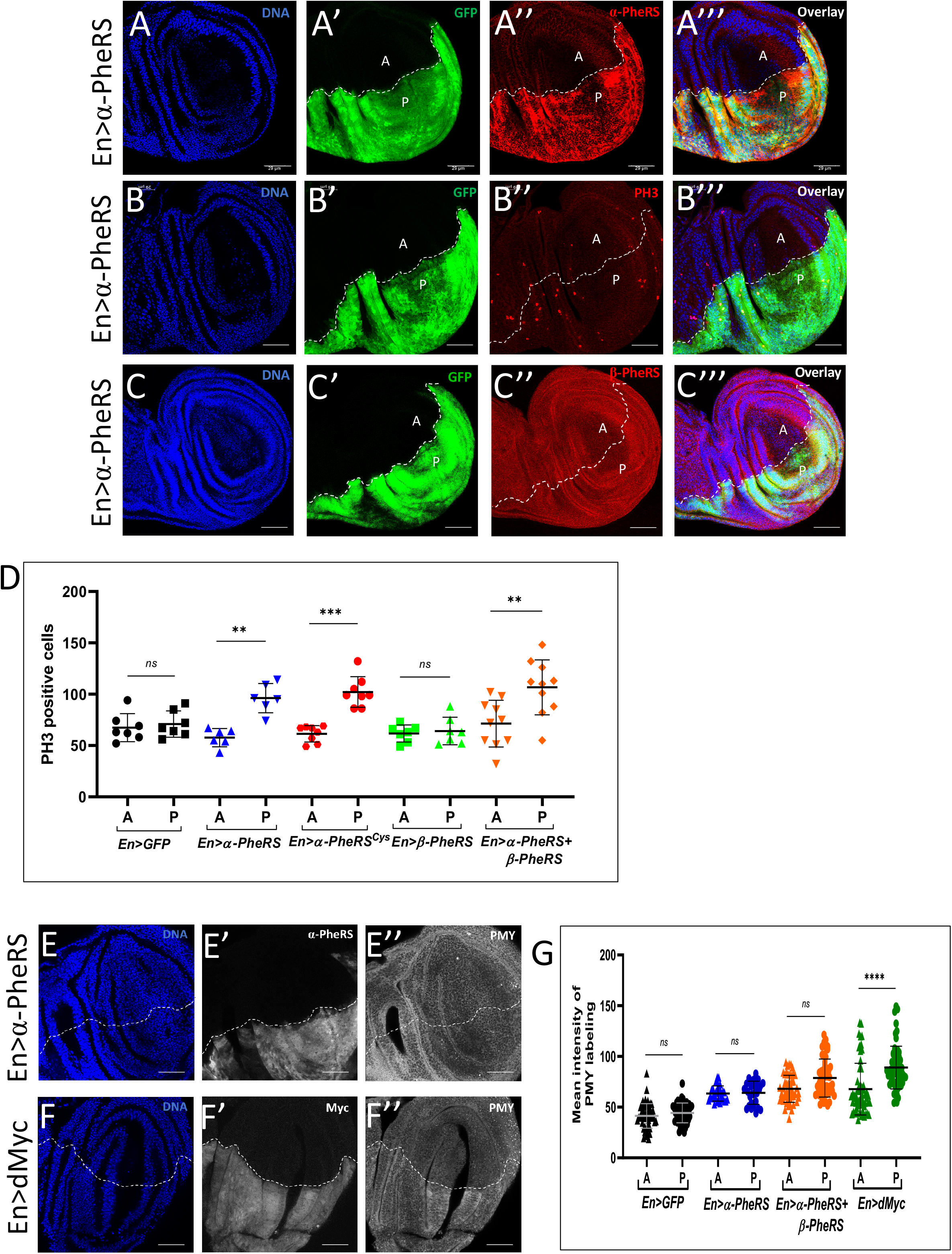
Elevated α–PheRS promotes the appearance of mitotic cells without *β-PheRS* and without stimulating translation. (A-C’’’,D) *en-Gal4/+;UAS-GFP/UAS-α-PheRS^(Cys)^* was used to drive transgene expression in order to elevate levels of *α-*PheRS or α-PheRS^Cys^ specifically in the posterior compartment of developing wing discs. (A”, A’’’) Upon staining with the anti-*α-*PheRS antibody, the average pixel intensity was 2-fold higher in the posterior compartment compared to the anterior one (n=5). (B”, B’’’) Mitotic cells were visualized with anti-phospho-Histone H3 (PH3) antibodies (A: anterior compartment; P: posterior compartment). n=10, *p<0.05, **p<0.01 in t-test. (C”) The increase of α-PheRS or α-PheRS^Cys^ levels did not affect the levels of β-PheRS. (E-F”, G) Protein synthesis did not increase upon overexpression of *α-PheRS* or *α-PheRS* and *β-PheRS* together. dMyc was used as a positive control (*en-Gal4/UAS-Myc::MYC;UAS-GFP/+*). *en-Gal4* was used to drive the overexpression of the transgenes in the posterior compartment of the wing discs. Protein synthesis was measured by the mean intensity of the puromycin (PMY) signal labeling the nascent polypeptides. n=15, ****p<0.0001, *ns:* not significant.

**Figure 4:**
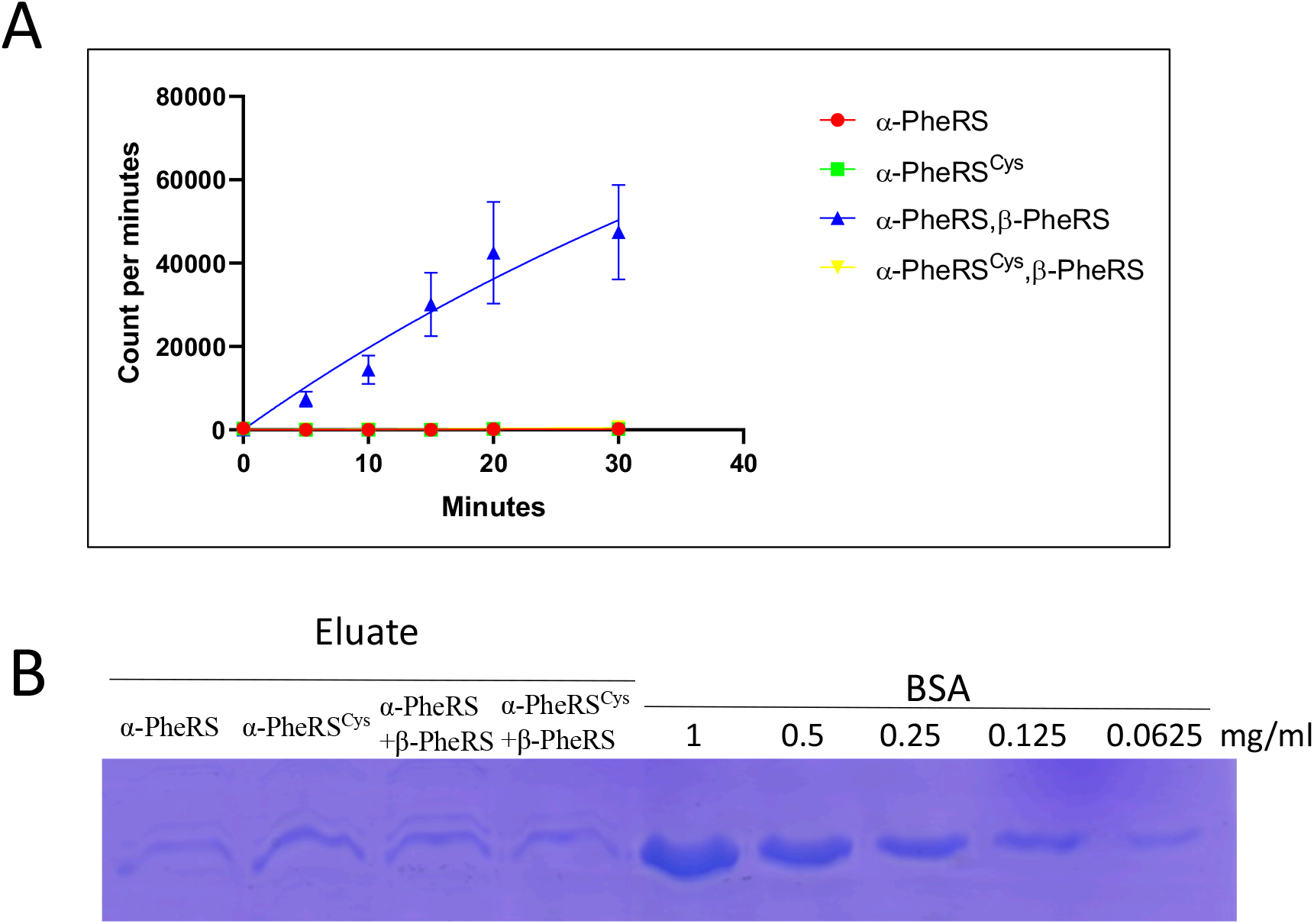
The *α–PheRS^Cys^* mutant does not support aminoacylation in vitro. (A) The aminoacylation assay was performed with the mixture of the recombinant protein (α–PheRS or α–PheRS^Cys^ and β-PheRS) simultaneously expressed in E. coli. tRNA^Phe^ from yeast was aminoacylated with [^3^H] phenylalanine. The [^3^H] phenylalanine decays were counted in a scintillation counter. CPM= counts per minute. In this assay, wild type α-PheRS + β-PheRS subunits together gave rise to CPM measurements between 38,000 and over 60,000 at 30 min. whereas α-PheRS^Cys^ + β-PheRS produced 350-820 CPMs, and α-PheRS or α-PheRS^Cys^ alone between 26 and 372 CPMs. (n=3) (B) Soluble recombinant proteins were affinity purified, eluted, and visualized by Coomassie staining, the protein amounts for each reaction were normalized by a BSA gradient.

### A non-canonical α-PheRS activity induces additional M-phase cells

To test directly whether α-PheRS can increase proliferation without stimulating translation, we made a mutant version of α-PheRS in which Tyr412 and Phe438 are replaced by cysteines (Cys). These substitutions are predicted to block the entrance into the phenylalanine binding pocket, preventing binding of Phe and aminoacylation of tRNA^Phe^ by the mutant PheRS^Cys^ (Finarov et al., 2010). To test whether the PheRS^Cys^ substitution indeed reduces the aminoacylation activity of PheRS, we expressed mutant and wild-type α-PheRS subunits individually or together with β-PheRS subunits in E. coli, His tag-purified them and performed aminoacylation assays with the same amounts of recombinant soluble proteins that were normalized to BSA standards (Fig 4B). The clear band in Fig 4B showed that the Cys mutation did not affect the solubility of the recombinant proteins. α-PheRS^Cys^ together with wild-type β-PheRS produced the same background signal as the α-PheRS subunit alone, and this background signal was only around 1% of the signal obtained with wild-type α-PheRS plus β-PheRS (Fig 4A). The Drosophila gene encoding the cytoplasmic α-PheRS subunit is located on the X chromosome. A P-element insertion into the 5’-untranslated region of the *α-PheRS* transcript (*α-PheRSP^2060^*) causes recessive lethality that can be rescued by a genomic copy of wild-type *α-PheRS* (*gα-PheRS*) that supports aminoacylation (Lu et al., 2014). A genomic copy of the aminoacylation defective *α-PheRS^Cys^, gα-PheRS^Cys^*, did not rescue the lethality of the *α-PheRSP^2060^* mutant, indicating that the Cys mutant is indeed not functional in aminoacylation *in vivo* in Drosophila. Despite this apparent lack of aminoacylation activity, expressing a transgenic copy of *α-PheRS^Cys^* (cDNA under UAS control) in the posterior compartment of the wing disc with the en-Gal4 driver caused a 67% increase in the number of mitotic cells in the above assay (Fig 3D). The fact that the mutant *α-PheRS^Cys^* version caused an increase in mitotic cells at least as strong as the wild-type *α-PheRS* expressed in the same way, together with the fact that β-PheRS overexpression was not needed for this effect, strongly suggests that the increased mitotic index is promoted without increasing the canonical function of PheRS.

We also tested directly whether elevated expression of wild-type α-*PheRS* and expression of *α-PheRS* and *β-PheRS* together are indeed unable to cause elevated translation as we expected. For this, we analyzed protein synthesis activity in the two wing compartments by puromycin (PMY) staining using the ribopuromycylation method (RPM) (Deliu et al., 2017). Testing this method, we first expressed elevated levels of the transcription factor dMyc in the posterior wing disc compartment. This positive control led to an increased protein synthesis activity and anti-PMY signal in the dMyc over-expressing posterior compartment relative to the anterior compartment of the same discs. In contrast, neither en-Gal4 driven expression of α-PheRS alone nor combined expression with β-PheRS increased the puromycin labeling in the posterior compartment (Fig 3E-F’’,G). The combined results therefore demonstrate unambiguously that elevated α-PheRS levels cause additional cells to be in mitosis through a non-canonical mechanism that does not involve a general increase in translation.

The non-canonical activity emanating from α-PheRS or α-PheRS^Cys^ is capable of inducing more cells to be in mitosis. Such a phenotype could come about by specifically slowing down progression through M-phase, causing higher numbers of cells to remain in the PH3-positive state. Alternatively, the activity might either promote the proliferation of mitotic cells or induce proliferation in non-cycling cells.

### *α-PheRS* levels accelerate growth and proliferation

To find out whether the cell cycle effect of elevated PheRS levels is a more general effect and to test the effect of PheRS levels on growth and proliferation directly, we set up “mosaic analysis with repressible cell marker” (MARCM; (Wu and Luo, 2006) assays in the ovarian follicle cells. Twin-spot clones were generated with one clone expressing elevated levels of PheRS and the GFP marker, and its twin clone expressing normal endogenous levels of PheRS and serving as an internal control (Fig 5A). The results of this experiment showed also in the follicle cells the PheRS levels affect the cell cycle because clonally elevated levels of both subunits of PheRS accelerated cell proliferation on average by 32% (Fig 5B). In contrast, increased expression of GFP with only the β-PheRS subunit or with GlyRS (also named GARS) did not significantly promote clonal expansion (Fig 5B). This shows that the stimulation of proliferation is specific for PheRS and not a general role of aaRSs. Interestingly, elevated expression of GFP with the α-PheRS subunit alone also stimulated cell proliferation autonomously by 30% (Fig 5B) and, intriguingly, this was very close to the 32% increase calculated for the clone overexpressing both PheRS subunits (Fig 5B). Remarkably, the higher number of mitotic cells observed upon α-PheRS overexpression in the posterior compartment of the larval wing discs (Fig 3C) was in a comparable range as the proliferation increase in the follicle cell assay (Fig 5B). These results, therefore, suggest that α-PheRS levels promote cell proliferation and that α-PheRS levels have this activity in different tissues.

**Figure 5:**
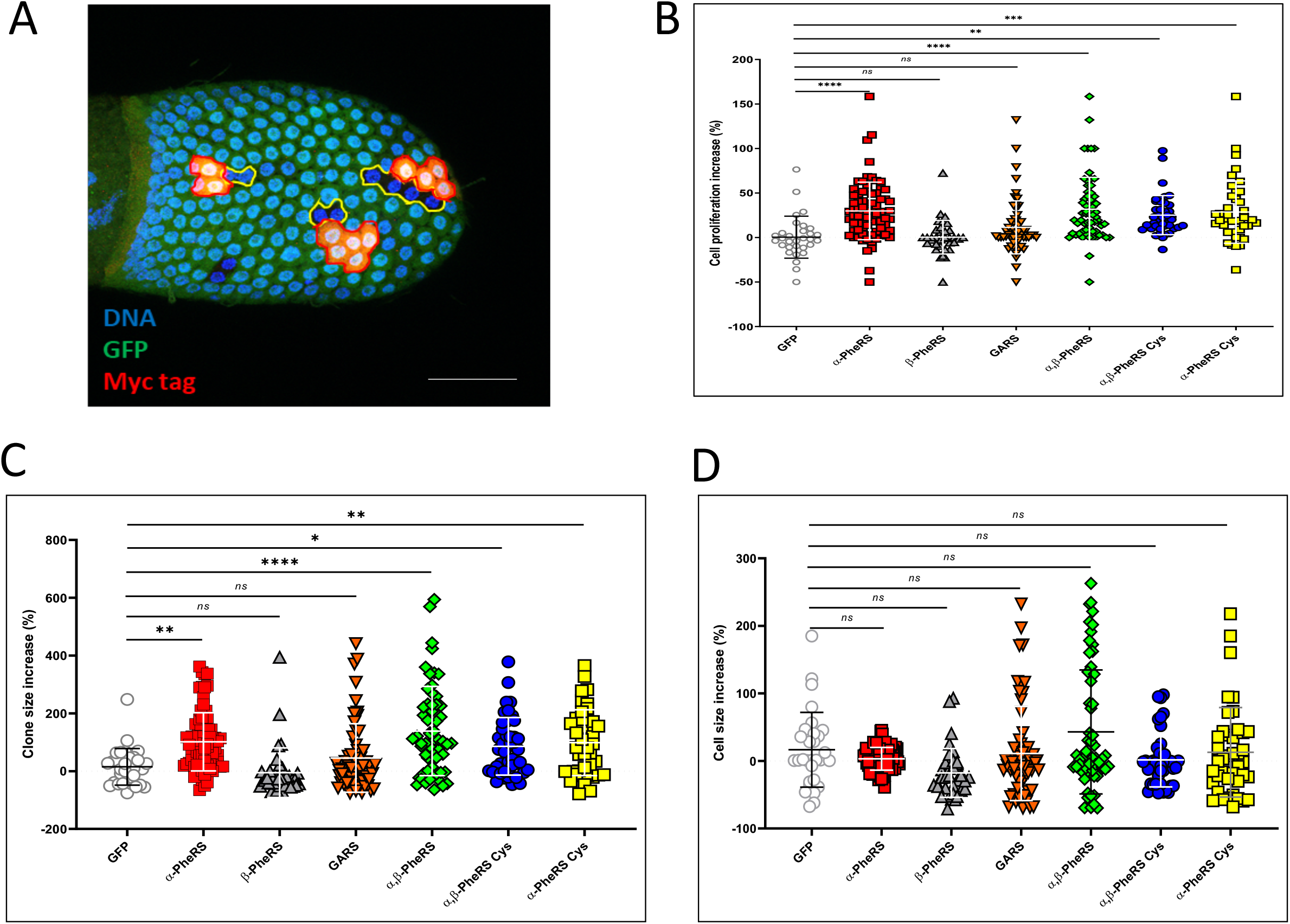
Aminoacylation-dead *α-PheRS* levels promote follicle cell growth and proliferation. Clonal twin spot analysis of the effect of overexpression of the PheRS subunits in follicle cells by the MARCM technique (*hs-flp; tub-Gal4/UAS-β-PheRS; FRT82B, ubiGFP, UAS-α-PheRS^(Cys)^,FRT82B Tub-Gal80*). Upon inducing mitotic recombination by a heat-shock (37°C for 40 minutes), a recombining follicle cell will divide and give rise to two proliferating clones. (A) One clone (red) overexpresses α-PheRS and also more GFP and its twin clone (GFP^-^) does not and expresses normal levels (internal control, outlined with yellow line). Note that for the presentation (but not for the quantification) a Myc-tagged α-PheRS was expressed to obtain a very clear picture. (B,C,D) Three days after inducing the recombination, cell numbers and clone sizes were determined and the average number of cell divisions and cell sizes were calculated for each clone and compared to its twin spot. (%). n= 30, **p<0.01, ***p<0.001, ****p<0.0001 in ANOVA test.

We also measured the clone size of the twin spots to find out whether the elevated *α-PheRS* could also enhance cell growth. Relative to the control twin spot, clonally elevated levels of both subunits of PheRS and α-PheRS subunits alone, respectively, augmented clone size by 139% and 102%, respectively. In contrast, elevated levels of single β-PheRS subunits or GlyRS did not show a significant increase in clone size (Fig 5C). We also observed that the size of single cells in twin spot clones remained unchanged (Fig 5D), indicating that cell size control was not affected and that cells overexpressing α-PheRS subunits also grow faster to sustain the higher proliferation. These results, therefore, show that α-PheRS levels promote cell growth and proliferation and they also suggest that α-PheRS levels have this activity in different tissues.

Elevated *α-PheRS^Cys^* alone and *α-PheRS^Cys^* together with β-*PheRS* (*PheRS^Cys^*) also stimulated cell proliferation in the follicle cell twin spot experiment by 28% and 25%, respectively (Fig 5B,C). Additionally, the clonal sizes also increased by 99% and 86%, respectively (Fig 5C). These observations confirmed that in addition to the proliferative activity, the stimulation of cell growth does also not depend on increased canonical PheRS activity.

## Discussion

Our work revealed that *PheRS* not only charges tRNAs with their cognate amino acid Phe but that it also performs a moonlighting function in stimulating cellular growth and proliferation. Because levels of α-PheRS are elevated in many tumor cells compared to their healthy counterparts (Park et al., 2019) and because a positive correlation between these levels and tumorigenic events had been noted some time ago (Sen et al., 1997), it was important to find out whether elevated PheRS levels are a mere consequence of the high metabolic activity of the tumor cells or whether they might also contribute to the over-proliferation of tumor cells. Here we now showed that α-PheRS has the potential to promote growth and proliferation and that it can do this independent of its aminoacylation activity, i.e. through a non-canonical or moonlighting activity. We found unambiguous evidence for the non-canonical nature of this proliferative activity and also showed that general translation is not elevated when this phenotype is induced by elevated α-PheRS or even the entire PheRS protein (Fig 3D-D”). Aminoacylation of tRNA^Phe^ requires both subunits to form the tetrameric protein α2β2-PheRS that can aminoacylate tRNA^Phe^ (Mosyak et al., 1995; Fig 4), and overexpression of an aminoacylation-dead *α-PheRS^Cys^* mutant subunit alone, (without simultaneous overexpression of the β-PheRS subunit) increased the cell numbers in the follicle cell clones as much as the wild-type gene did when expressed in the same way (Fig 4, 5B). The described non-canonical functions of α-PheRS are not only independent of the aminoacylation activity, they also do not reflect a function in sensing the availability of its enzymatic substrate, Phe, for the major growth controller, the TOR kinase (Fig 2). Such activities have been described for other members of the aaRS family, TrpRS or LeuRS (Adam et al., 2018; Bonfils et al., 2012a; Han et al., 2012a).

The notion that the α-PheRS subunit can be stable (Fig 3A”) and function independently of the β-subunit (Fig 3C-C’”) was surprising because previous results showed that the two subunits were dependent on the presence of the other subunit for their stability (Antonellis et al., 2018; Lu et al., 2014; Xu et al., 2018). Our results now show that this requirement does not apply to all cell types. In mitotically active follicle cells, the posterior compartment of the wing disc and possibly other cells, elevated expression of the α-PheRS subunit alone produced a strong cell cycle phenotype. This suggests that the α- and β-PheRS subunits function together in every cell to aminoacylate tRNA^Phe^, but in addition, the α-subunit can be stable in specific cell types that appear to have retained their mitotic potential. In these cells, α-PheRS assumes a novel function in promoting cell growth and proliferation. This would then suggest that many differentiating cell types start to put a system in place that prevents α-PheRS accumulation at high levels. Such a mechanism could then contribute to reducing the proliferative activity of differentiated cells.

PheRS, TrpRS, and LeuRS are not the only aaRS family member with roles beyond charging tRNAs (Dolde et al., 2014; Guo et al., 2010; Lu et al., 2015; Pang et al., 2014). For instance, MetRS/MRS is capable of stimulating rRNA synthesis (Ko et al., 2000), GlnRS/QRS can block the kinase activity of apoptosis signal-regulating kinase 1 (ASK1) (Ko et al., 2001) and a proteolytically processed form of YARS/TyrRS acts as a cytokine (Casas-Tinto et al., 2015; Greenberg et al., 2008). aaRSs are, however, also not the only protein family which evolved to carry out more than one function. In recent years, it has become increasingly evident that many, if not most, proteins have evolved to carry out not only one, but two or more functions, providing interesting challenges to figure out, which of their activities are important for the specific function of a gene (Dolde et al., 2014).

Improper expression of PheRS was suspected long ago to promote carcinogenesis, but till now the mechanisms behind this effect remained unknown. Elevated mRNA levels of the human *α-PheRS, FARSA*, during the development of myeloid leukemia correlate with tumorigenic events and many malignant cell lines express elevated levels of mRNA encoding the PheRS subunits compared to normal human tissue (Sen et al., 1997; Rodova et al., 1999). The GENT2 database published in 2019 describes also strong positive correlations between PheRS subunit mRNA levels and tumorigenic events in several tissues and cancers (Barker et al., 2009; Park et al., 2019). Interestingly, and consistent with our results, not all tumors that displayed elevated PheRS levels showed elevated levels of *α*- and *β-PheRS* mRNA. For instance, brain, ovary, endometrium, and bladder tumors displayed only elevated *α-PheRS* mRNA levels while colon, breast, lung, and liver tumors showed elevated levels of mRNAs for both subunits. Because elevated levels of *α-PheRS* or *α-PheRS^Cys^* alone can elicit mitotic activity, growth, and proliferation, our results suggest that the excessive PheRS (FARS) levels in tumor tissues might be able to produce such proliferative signals independent of whether they also produce elevated levels of β-PheRS. Modeling the effect of elevated α-PheRS levels in *Drosophila*, we found that elevated levels support growth and proliferation and lead to an increase in mitotic cells in different cell types. In follicle cells, more cells were produced in clones expressing more α-PheRS compared to wild-type clones. In wing discs, more mitotic cells were detected in most areas with higher levels of α-PheRS. This indicates that elevated α-PheRS levels can indeed be a risk factor for tumor formation in several different tissues.

## Materials and Methods

### Fly genetics and husbandry

All *Drosophila melanogaster* fly stocks were kept for long term storage at 18°C in glass or plastic vials on standard food with day/night (12h/12h) light cycles. All experiments were performed at 25°C unless specifically mentioned. A UAS-GFP element was added in experiments that tested for rescue and involved Gal4 mediated expression of the rescue gene. This construct served to even out the number of UAS sites in each Gal4 expressing cell. Origins of all stocks are noted in Table 1, the *Key Resource Table*.

**Table 1:**
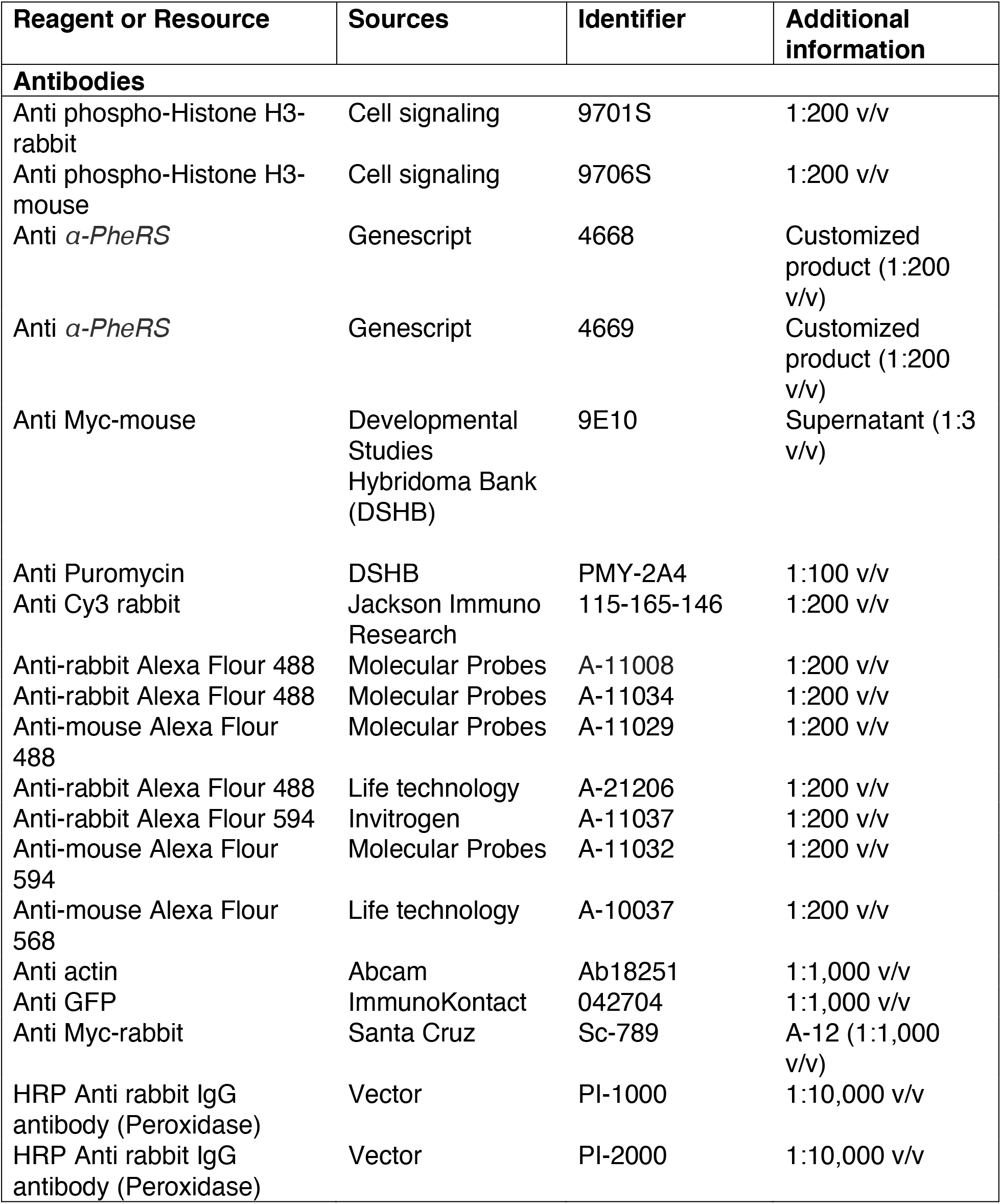
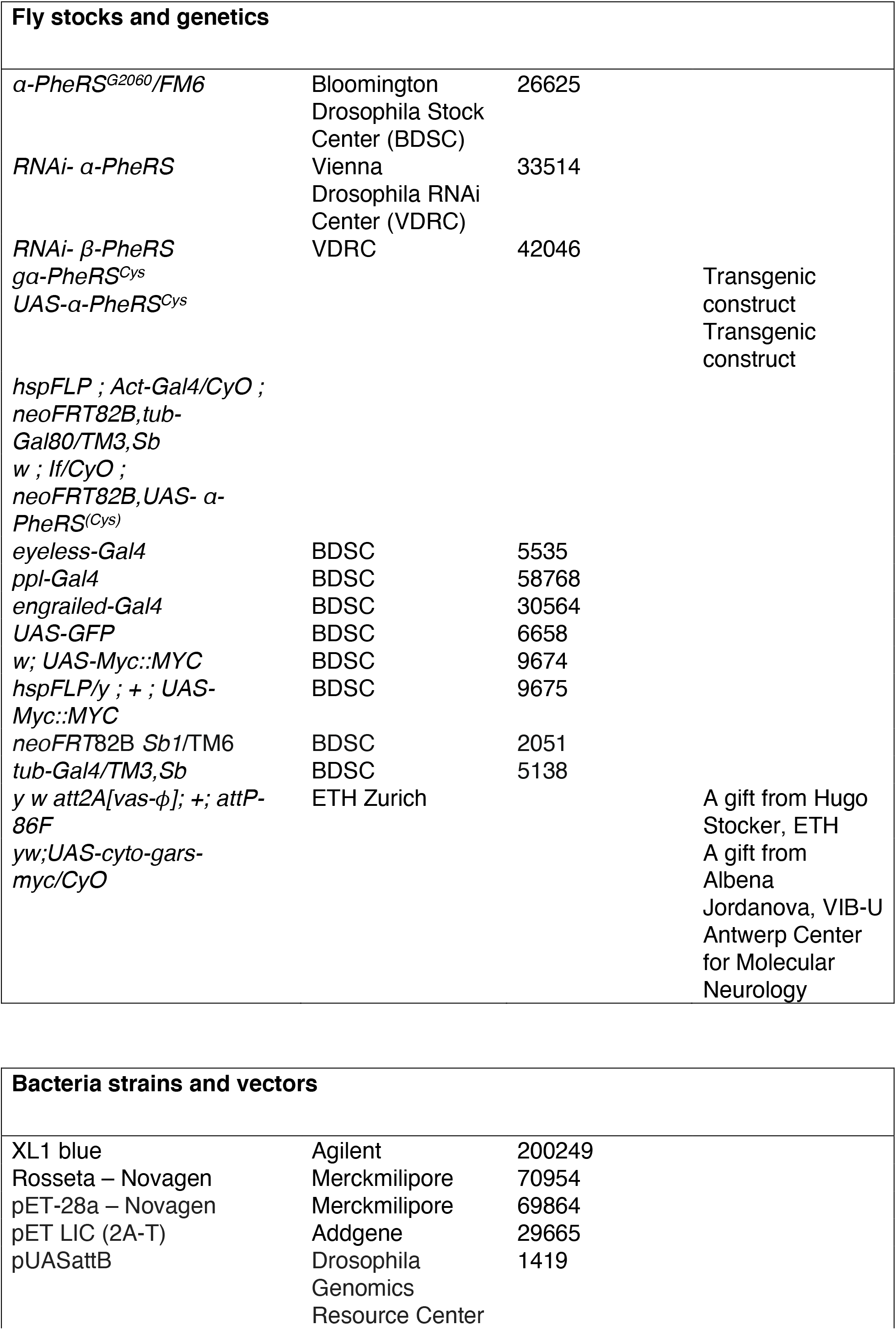
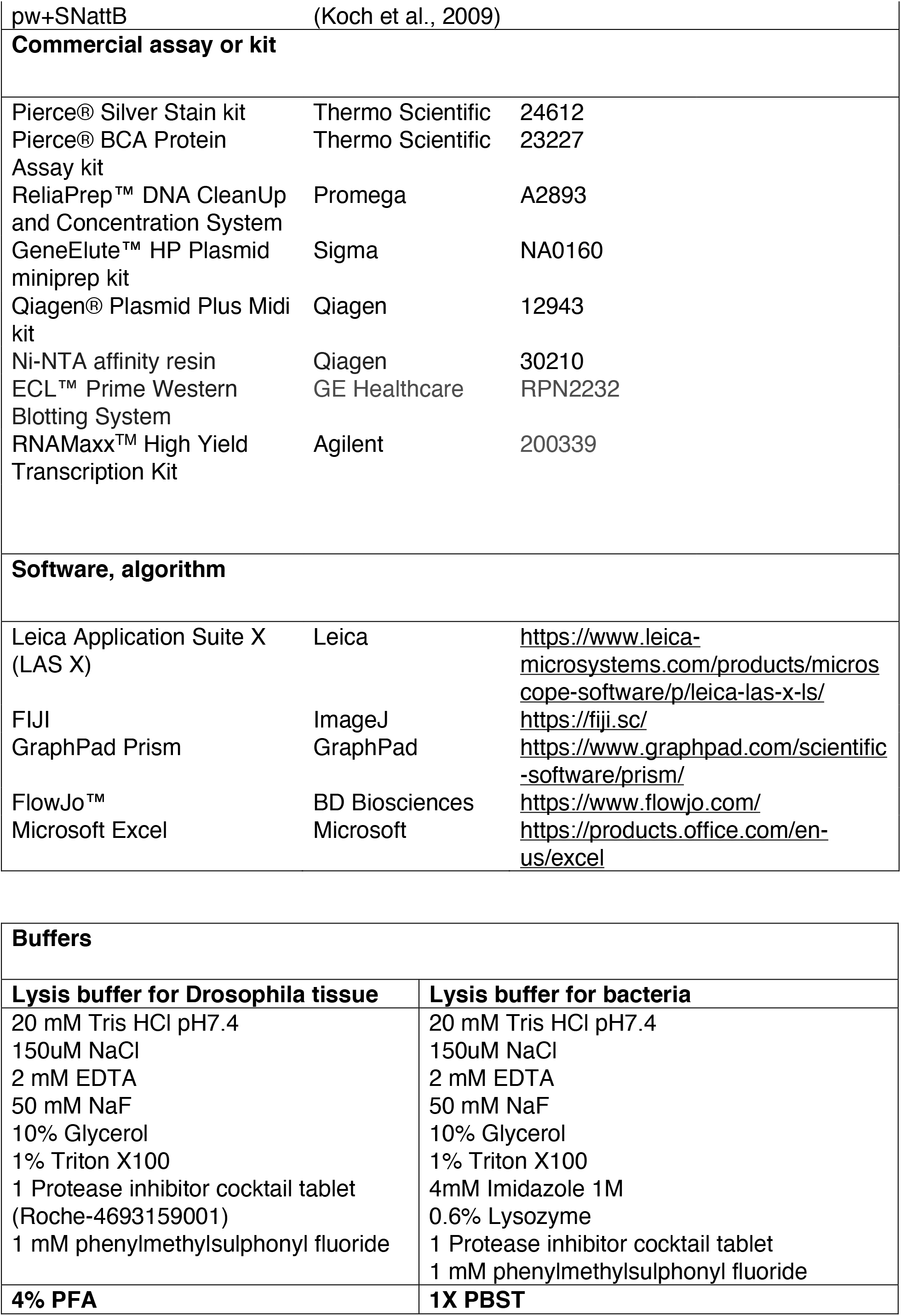
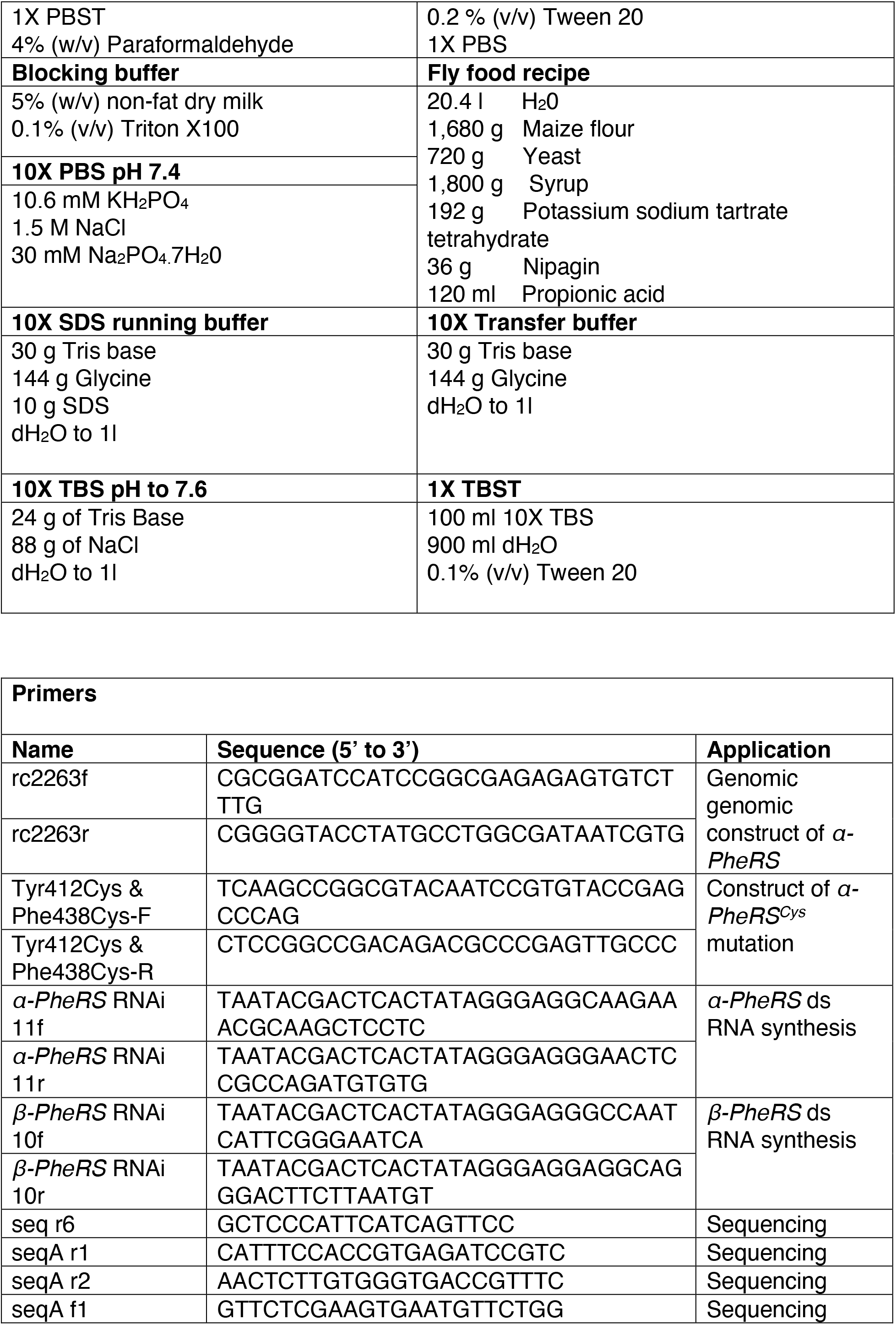
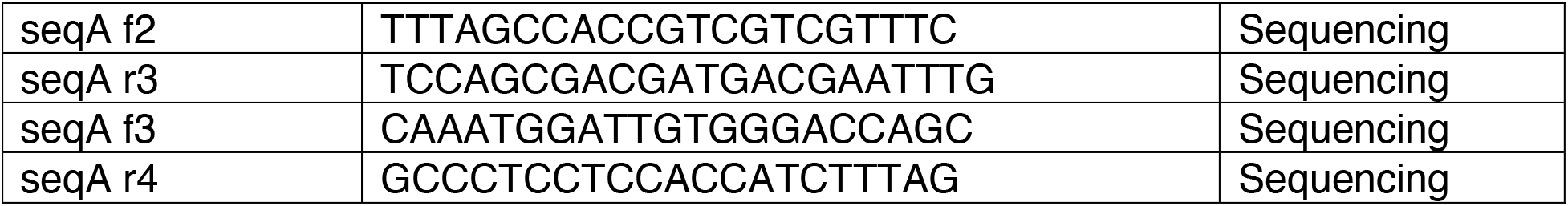
Key Resources Table

### DNA cloning and generation of transgenic flies

Sequence information was obtained from Flybase. All mutations and the addition of the Myc-tag to the N-terminus of *α-PheRS* were made by following the procedure of the QuickChange^®^ Site-Directed Mutagenesis Kit (Stratagene). The genomic *α-PheRS* rescue construct (*Myc::α-PheRS*) codes for the entire coding region and for an additional Myc tag at the N-terminal end. In addition, it contains ~ 1kb of up- and down-stream sequences and it was cloned into the *pw^+^SNattB* transformation vector (Koch et al., 2009; Lu et al., 2014). The *α-PheRS* and *β-PheRS* cDNAs were obtained by RT-PCR from mRNA isolated from 4-8 days old *OreR* flies (Lu et al., 2014). The Tyr412Cys and Phe438Cys mutations in the *α-PheRS* sequence were created by site directed mutagenesis. Like the wild-type cDNA, they were cloned into the *pUASTattB* transformation vector to generate the pUAS-α-PheRS and pUAS-α-PheRS^Cys^. Before injecting these constructs into fly embryos, all plasmids were verified by sequencing (Microsynth AG, Switzerland). Transgenic flies were generated by applying the *φ* C31-based integration system with the stock (*y w att2A[vas-ϕ]; +; attP-86F*) (Bischof et al., 2007).

### Western blotting

Protein was extracted from tissues, whole larvae, or flies using lysis buffer. Protein lysates were separated by SDS-PAGE and transferred onto PVDF membranes (Milipore, US). The blocking was performed for 1h at room temperature (RT) with non-fat dry milk (5%) in TBST solution. Blots were probed first with primary antibodies (diluted in blocking buffer) overnight at 4°C and then with secondary antibodies (diluted in TBST) 1h at RT. The signal of the secondary antibody was detected by using the detect solution mixture (1:1) (ECL™ Prime Western Blotting System, GE Healthcare Life Science) and a luminescent detector (Amersham Imager 600, GE Healthcare Life Science). Origins and recipes of all buffers and reagents are noted in Table 1, the *Key Resource Table*.

### Immunofluorescent staining and confocal microscopy

Dissections were performed in 1x PBS on ice and tissues were collected within maximum one hour. Fixation with 4% PFA in PBS-T 0.2% at RT was done for 30 minutes (wing discs, ovaries). Then the samples were blocked overnight with blocking buffer at 4°C. Primary antibodies (diluted in blocking buffer) were incubated with the samples for 8h at RT. The samples were rinsed 3 times and washed 3 times (20 minutes/wash) with PBST. Secondary antibodies (diluted in PBST) were incubated overnight at 4°C. The samples were then rinsed 3 times and washed 2 times (20 minutes/wash) with PBST. Hoechst 33258 (2.5 μg/ml) was added in PBST before the last washing step and the samples were mounted with Aqua/Poly Mount solution (Polysciences Inc., US). Origins and diluted concentrations of all buffers and antibodies are noted in Table 1, the *Key Resource Table*.

### Protein synthesis measurements using the ribopuromycylation method (RPM)

For puromycin labeling experiments, tissues were dissected in Schneider’s insect medium (Sigma, US) supplement with 10% fetal calf serum (FCS, Sigma, US) at 25°C. They were then incubated with Schneider’s insect medium containing puromycin (5 μg/ml, Sigma, US) and cycloheximine (CHX, 100 μg/ml, Sigma, US) for 2 hours at RT. Then the samples were fixed with 4% PFA in PBS-T 0.2% at RT and blocked overnight with blocking buffer at 4°C. Primary anti-Puromycin antibody (diluted in PBST) was incubated with the samples for 8h at RT. The samples were rinsed 3 times and washed 3 times (20 minutes/wash) with PBST. Secondary antibodies (diluted in PBST) were incubated overnight at 4°C. The samples were then rinsed 3 times and washed 2 times (20 minutes/wash) with PBST. Hoechst 33258 (2.5 μg/ml) was added in PBST before the last washing step and the samples were mounted with Aqua/Poly Mount solution (Polysciences Inc., US).

### *In vitro* aminoacylation assay

Recombinant α-PheRS and β-PheRS proteins were co-expressed in the *E. coli* strain Rosetta (Novagen) and then purified (Moor et al., 2002). For this, the *α-PheRS* or *α-PheRS^Cys^* mutant cDNAs were cloned with His tags at the N-terminal end into the pET-28a plasmid expression vector (Novagen). Wild-type *β-PheRS* cDNAs were cloned into the pET LIC (2A-T) plasmid (Addgene). Then, His-α-PheRS or the His-α-PheRS^Cys^ mutant and β-PheRS were co-expressed in the *E. coli* strain Rosetta with isopropylthiogalactoside (IPTG, 1mM) induction at 25 °C for 6 hours. Proteins were purified with Ni-NTA affinity resin (Qiagen). The aminoacylation assay protocol from Jiongming Lu was then followed (Lu et al., 2014) with the modification that the Whatman filter paper discs were soaked in Phenylalanine solution for 1 hour (30mg/ml in 5% trichloroacetic acid (TCA)) to reduce the background. This assay was performed at 25 °C in a 100μl reaction mixture containing 50 mM Tris-HCl pH 7.5, 10 mM MgCl_2_, 4 mM ATP, 5 mM β-mercaptoethanol, 100 μg/ml BSA, 3 U/ml *E. coli* carrier tRNA, 5 μM [^3^H]-amino acid (L-Phe) and 1 μM tRNA^Phe^ from brewer’s yeast (Sigma, US). In each experiment, a 15-μl aliquot was removed at four different incubation time points, spotted on the Phe-treated Whatman filter paper discs and washed three times with ice-cold 5% TCA and once with ice-cold ethanol. A blank paper disc without spotting and another with spotting the enzyme-free reaction were used for detecting background signals. After filter discs were dried, they were immersed in PPO Toluol (Sigma, US) solution in plastic bottles and the radioactivity was measured by scintillation counting.

### Wing disc dissociation and FACS analysis

Wandering larvae derived from 2-4 hours egg collections were dissected in PBS during a maximal time of 30 minutes. Around twenty wing discs were incubated with gentle agitation at 29°C for around 2-hours in 500μl 10× Trypsin-EDTA supplemented with 50 μl 10×Hank’s Balanced Salt Solution (HBSS) (Sigma, US) and 10 μl Vybrant DyeCycle Ruby stain (Molecular Probes, US). Dissociated cells from wing discs were directly analyzed by FACS-Calibur flow cytometer (Becton Dickinson, US).

*Drosophila* tissue culture cells were harvested and fixed in 70% ethanol and stained with a staining solution containing 1mg/ml propidium iodide, 0.1% Triton and 10 mg/ml RNase A. The cells were then subjected to FACS-Calibur cytometry and data were analyzed with the FlowJo software.

### *Drosophila* cell culture and RNAi treatment

*Drosophila* Kc cells were incubated at 25°C in Schneider’s *Drosophila* medium supplemented with 10% heat-inactivated fetal calf serum (FCS) and 50 μg/ml Penicillin/Streptomycin. To induce RNAi knockdown in *Drosophila* cells, dsRNA treatment was performed (Clemens et al., 2000). dsRNAs around 500bp in length were generated with the RNAMaxx™ High Yield Transcription Kit (Agilent, US). Cells were diluted to a concentration of 10^6^ cells/ml in serum-free medium, and dsRNA was added directly to the medium at a concentration of 15 μg/ml. The cells were incubated for 1 hour followed by addition of medium containing FCS. Then the cells were kept in the incubator and were harvested at different time points 1-5 days after dsRNA treatment.

### Clonal assay and twin spot data analysis

For twin spot tests, we used the Mosaic Analysis with a Repressible Cell Marker (MARCM) system. Twin spots were generated with the progenitor genotype *hs-flp; tub-Gal4/UAS-β-PheRS; FRT82B, ubiGFP, UAS-α-PheRS^(Cys)^/FRT82B Tub-Gal80*. In twin spots, the internal control clone was GFP-minus and the twin sister clone produced a red signal by the antibody against the overexpressed protein. We induced the *hs-FLP, FRT82B* system at 37°C for 1h on the third day post-eclosure and dissected the animals 3 days post-induction. Confocal imaging detected non-green clones (without ubiGFP) and red clones (stained with Myc antibody-red) (Fig 5A).

In twin spots, cell numbers per clone were counted and the numbers of cell divisions per clone were calculated as log_2_(cell numbers per clone). This represents the logarithm of the cell numbers per clone to the base 2. The increase of cell proliferation (%) was analyzed by comparing the number of cell divisions of the clone pairs in the same twin spot. The clone sizes were measured by FIJI software and the increase of clone size was analyzed by comparing the clone size in the same twin spot.

### Image acquisition and processing

Imaging was carried out with a Leica SP8 confocal laser scanning microscope equipped with a 405 nm diode laser, a 458, 476, 488, 496 and 514 nm Argon laser, a 561 nm diode pumped solid state laser and a 633 nm HeNe laser. Images were obtained with 20x dry and 63x oil-immersion objectives and 1024×1024 pixel format. Images were acquired using LAS X software. The images of the entire gut were obtained by imaging at the standard size and then merging maximal projections of Z-stacks with the Tiles Scan tool. Fluorescent intensity was determined from FIJI software.

### Quantification and statistical analysis

For quantifications of all experiments, *n* represents the number of independent biological samples analyzed (the number of wing discs, the number of twin spots), error bars represent standard deviation (SD). Statistical significance was determined using the t-test or ANOVA as noted in the figure legends. They were expressed as P values. (*) denotes p < 0.05, (**) denotes p < 0.01, (***) denotes p < 0.001, (****) denotes p < 0.0001. (*ns*) denotes values whose difference was not significant.

## Acknowledgements

We thank Hugo Stocker, Albena Jordanova, Erik Storkebaum, and the Bloomington Stock Center for fly stocks. We are also grateful to Mark Safro for suggesting mutations that disrupt the phenylalanine binding site of α-PheRS. We further thank Dominique Brunssen and Jonathan Huot for their valuable advice on the aminoacylation assay. This work was supported by the Novartis Foundation for medical-biological Research (#18A050), the Swiss National Fund (project grant 31003A_173188), and the University/Canton of Bern.

